# ClC-2 contributes to hypotonicity-induced adrenal aldosterone secretion

**DOI:** 10.1101/2025.03.19.644253

**Authors:** Marina Volkert, Hoang An Dinh, Ute I. Scholl, Gabriel Stölting

## Abstract

The zona glomerulosa (ZG) of the adrenal cortex regulates blood pressure and electrolyte homeostasis through aldosterone production. In ZG cells, the serum concentrations of potassium and angiotensin II (Ang II) trigger calcium oscillations that drive aldosterone synthesis. Changes in serum osmolality also modulate aldosterone production in a chloride-dependent fashion, but the involved proteins remain unclear.

Because the chloride channel ClC-2 is activated by hypoosmolality, we investigated its role in ZG osmoregulation using ClC-2 knockout (KO) mice. Intracellular chloride concentrations in the ZG are high, and opening of ClC-2 leads to chloride efflux, depolarization and voltage-dependent calcium influx.

Under hypoosmolar conditions, intracellular chloride levels were higher in ClC-2 KO ZG cells than in the WT, suggesting that hypoosmolality triggers chloride efflux via ClC-2 in the WT, and that this efflux is absent in the KO. WT cells responded to hypoosmolality with an increase in intracellular calcium levels, likely mediated by chloride efflux and depolarization. This response was again abrogated in the KO, despite faster calcium spiking. In line with increased intracellular calcium levels, WT adrenal slices upregulated aldosterone production upon hypoosmolar treatment in vitro, whereas aldosterone production remained unchanged in the KO.

These findings establish a role for ClC-2 in the ZG’s response to reductions in extracellular osmolality through the outflow of chloride, voltage-dependent calcium influx and aldosterone production, advancing our general understanding of regulators of aldosterone production and the specific role of ClC-2.

## Introduction

The adrenal glands, paired organs located cranial to the kidneys, consist of the neuroendocrine medulla as well as the steroid-hormone producing cortex. The cortex comprises distinct functional layers, with the zona glomerulosa (ZG) forming the outermost layer responsible for mineralocorticoid synthesis. The mineralocorticoid aldosterone is required for maintaining salt and water homeostasis, acting primarily via altering water and salt reabsorption in the kidneys and intestine.

Because aldosterone is lipophilic, ZG cells regulate its release at the level of production from cholesterol, not via exocytosis. The primary stimuli are the serum concentrations of angiotensin II (Ang II) and potassium^1^. Under physiological conditions, Ang II triggers membrane depolarization through the closure of background potassium channels, leading to a depolarization of the ZG^2^, whereas hyperkalemia directly depolarizes the cell. This facilitates calcium influx via T- and L-type voltage gated calcium channels^3^, regulating aldosterone synthesis^4^.

Mutations in several ion channel genes enhance this depolarization or calcium influx, resulting in an excessive and uncontrolled synthesis of aldosterone in a disease termed primary aldosteronism^5–8^. Among these, gain-of-function variants in the voltage-gated chloride channel ClC-2 (encoded by the *CLCN2* gene) were identified in the germline of patients with familial hyperaldosteronism^9,10^ but also as somatic mutations in aldosterone-producing adenomas^11^.

ClC-2 is a nearly ubiquitously expressed, inwardly rectifying chloride channel^12,13^. Apart from its role in primary aldosteronism^14^, various other pathologies are associated with gain- or loss-of-function variants, including in brain^15,16^ and heart^17^.

Despite the thorough investigations into the effects of pathogenic variants, we know much less about the physiological role of ClC-2 in mammalian cells and tissues. The best evidence for a physiological role of ClC-2 exists for the murine colonic epithelium where the basolateral channel mediates chloride reabsorption^18,19^. ClC-2 is also important for Sertoli cells in testes and the retinal pigment epithelium where its loss leads to testicular^20^ and retinal degeneration^20–22^, although the precise molecular mechanisms remain unclear^23^.

ZG cells possess a high intracellular chloride concentration of about 70 mmol/l^9^. ClC-2 activates upon voltages negative to the equilibrium potential for chloride. Gain-of-function variants in ClC-2 therefore result in a depolarizing chloride efflux that leads to higher voltage-dependent calcium influx and increased aldosterone synthesis^24,25^. Unlike in aldosteronism-linked mutations in voltage-gated calcium channels^26,27^, the calcium influx is the result of an increased frequency of calcium oscillations^14,24,25^. This, however, does not explain the physiological role of wild-type ClC-2 in the ZG. We initially hypothesized that its slow hyperpolarization-dependent activation might mediate the initial depolarization required for voltage oscillations in the ZG^9,24^ but ClC-2 KO mice show normal aldosterone levels^24,25^, arguing against such a role.

There is a longstanding debate on the physiological role of ClC-2’s activation by hypoosmolality^12,28^, and it has been implicated in osmoregulation with the best examples being in heterologous expression systems^28^ and in malaria infected cells^29^.

Given the known chloride-dependent osmo-sensitivity of the ZG^30–32^, we set out to examine the involvement of ClC-2. For this, we investigated acute slice preparations of murine WT and ClC-2 KO adrenal glands using chloride and calcium imaging.

## Results

### ClC-2 mediates chloride efflux under hypoosmolar conditions

To test whether the activation of ClC-2 by hypoosmolality leads to the efflux of chloride, we measured intracellular chloride concentrations in the murine zona glomerulosa using the fluorescent indicator MEQ (Fig. 1A).

**Figure 1.**
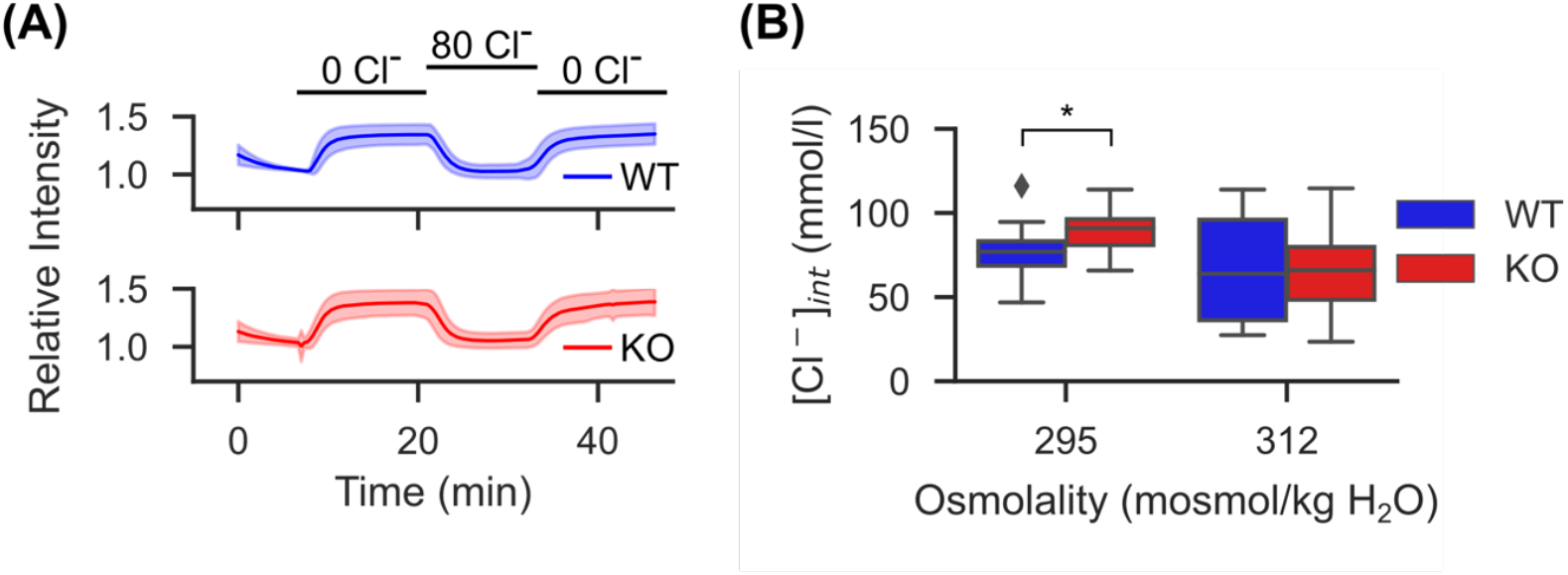
Extracellular hypoosmolality leads to increases of intracellular chloride in *Clcn2* KO mice. **(A)** Average (thick line) ± SD (light area) of MEQ fluorescence traces at 295 mosmol/kg H_2_O. Initially, standard extracellular solution was used and then exchanged for calibration solutions containing 0 or 80 mM Cl^−^ as indicated. All traces were individually normalized to the value prior to the first perfusion with 0 mM Cl^−^ for display. Steady-state values at the end of each perfusion condition were taken as representative of the chloride-dependent fluorescence for the indicated concentration. **(B)** Chloride concentrations for the genotypes and conditions indicated. (All values mean ± SD: @312 mosmol/kg H_2_O, WT: 65.5 ± 29.1 mM, n = 17 recordings; KO: 68.0 ± 26.5 mM, n = 13 recordings; p = 0.81, t = −0.23; @295 mosmol/kg H_2_O: WT: 75.6 ± 16.9 mM, n = 15 recordings; KO: 89.3 ± 12.8 mM, n = 16 recordings p = 0.019, *, t = −2.48; Student’s t-test). Box plots are shown as defined in Materials & Methods.

Mice exhibit higher isotonicity than humans, which is also strain dependent^33,34^. Published values for C57BL/6 mice average at ∼312 mosmol/kg H_2_O, which we chose as isotonic for this study. Under such conditions, both genotypes showed similar baseline chloride concentrations (Fig. 1B). In contrast, during hypotonic exposure (295 mosmol/kg H_2_O), KO cells presented higher chloride concentrations than WT cells, demonstrating a role of ClC-2 in the lowering of intracellular chloride levels during phases of hypotonicity (Fig 1B).

### Loss of ClC-2 prevents osmolality-induced increases of intracellular calcium

The activation of a chloride efflux pathway should lead to a depolarization of ZG cells, facilitating the influx of calcium via the activation of voltage-gated calcium channels, elevating intracellular calcium concentrations. Using ratiometric Fura-2 imaging to measure absolute calcium concentrations, we found that WT ZG cells indeed showed significantly higher intracellular calcium concentrations when exposed to hypotonic solutions for different concentrations of Ang II (Fig. 2A). This effect was less prominent in the presence of high extracellular K^+^ concentrations (Fig. 2B).

**Figure 2.**
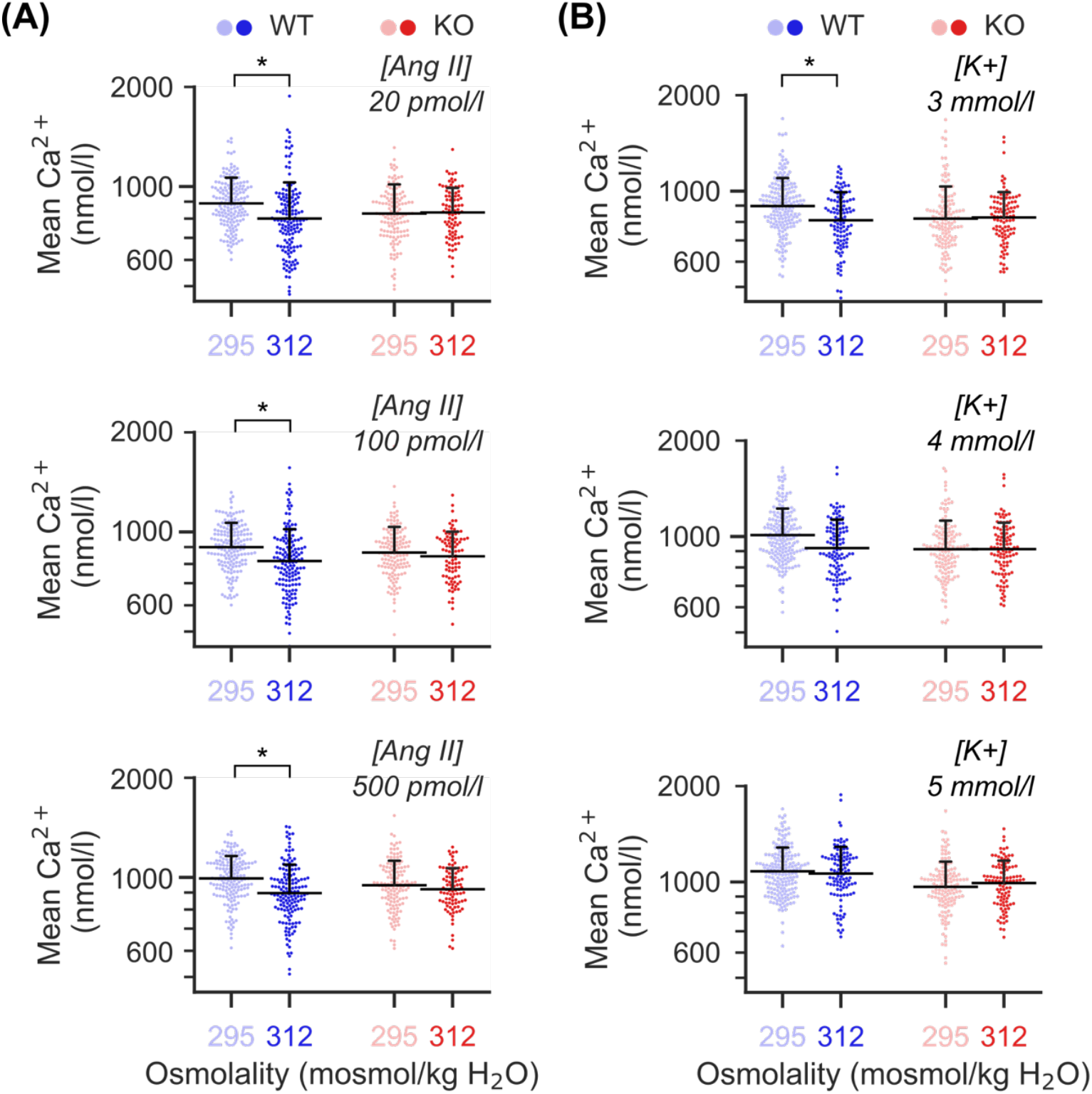
Absolute intracellular calcium concentrations increase under some hypotonic conditions in WT but not KO. **(A)** Absolute intracellular calcium concentrations at three different stimulations with Ang II (shown in italic font) and either hypotonic (295 mosmol/kg H_2_O, light colors) or isotonic conditions (312 mosmol/kg H_2_O, dark colors). Extracellular K^+^ was kept constant at 4 mmol/l. Absolute [Ca^2+^] values were determined following calibration of 340/385 nm Fura-2 signals as described^26^. In the ZG from WT mice (blue), the intracellular calcium concentration increased with hypotonicity while it remained unaffected in cells from KO mice (red). (p_WT,20AngII_ = 0.02, *; p_WT,100AngII_ = 0.03, *; p_WT,500AngII_ = 0.01, *; p_KO,20AngII_ = 0.59; p_KO,100AngII_ = 0.51; p_KO,500AngII_ = 0.58; likelihood ratio tests of linear mixed models; further information in Supp. Table 1). Each dot represents one cell, horizontal black lines indicate the mean, with the whisker representing the SD (only the positive whisker is shown for clarity). **(B)** Absolute intracellular calcium concentrations at three different concentrations of extracellular K^+^ (shown in italic font). Ang II was kept constant at 100 pmol/l. In the ZG from WT mice (blue), the intracellular calcium concentration only increased significantly at 3 mmol/l of K^+^ and remained unaffected in cells from KO mice (red). (p_WT,3K+_ = 0.03, *; p_WT,4K+_ = 0.07; p_WT,5K+_ = 0.46; p_KO,3K+_ = 0.98; p_KO, 4K+_= 0.82; p_KO, 5K+_= 0.66; likelihood ratio test of linear mixed model; further information in Supp. Table 2). Each dot represents one cell, horizontal black lines indicate the mean, with the whisker representing the SD (only the positive whisker is shown for clarity).

In contrast, KO cells maintained baseline calcium levels and did not respond to changes in extracellular osmolality regardless of stimulation with Ang II (Fig. 2A) or K^+^ (Fig. 2B).

**Table 1:**
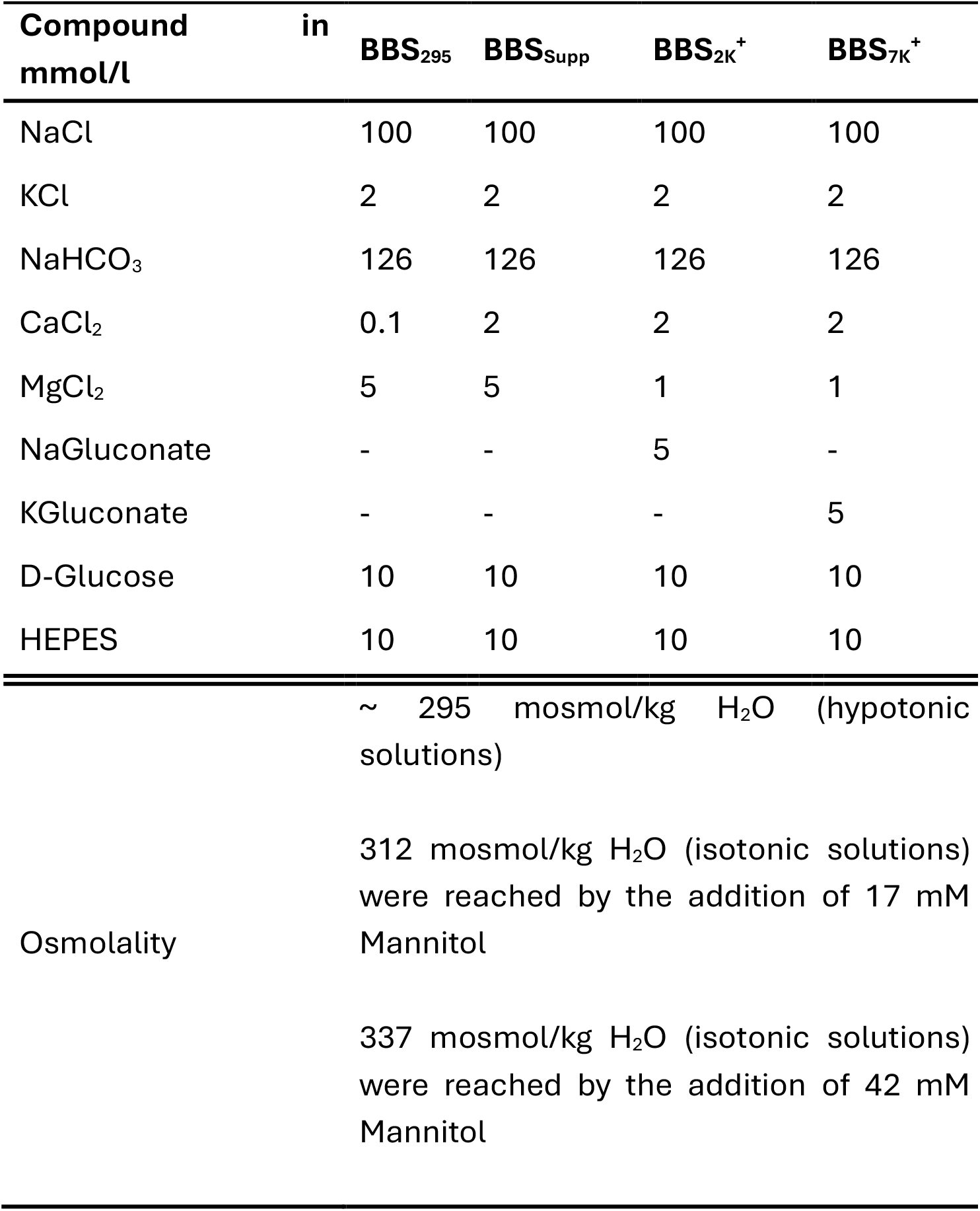
Composition of BBS solutions.

### Hypoosmolality increases aldosterone production in WT but not *Clcn2* KO mice

To test whether the increase in intracellular calcium levels also led to higher aldosterone synthesis, we incubated adrenal glands in vitro. We extracted both adrenal glands from either WT or KO mice. Each collected gland was incubated separately. Initially, they were both kept at 312 mosmol/kg H_2_O (with continuous oxygen supply). Then, we changed the solution for one adrenal gland to an osmolality of 295 mosmol/kg H_2_O while the other one remained in fresh 312 mosmol/kg H_2_O solution. Aldosterone levels were determined after a total of 120 minutes (Fig. 3).

**Figure 3.**
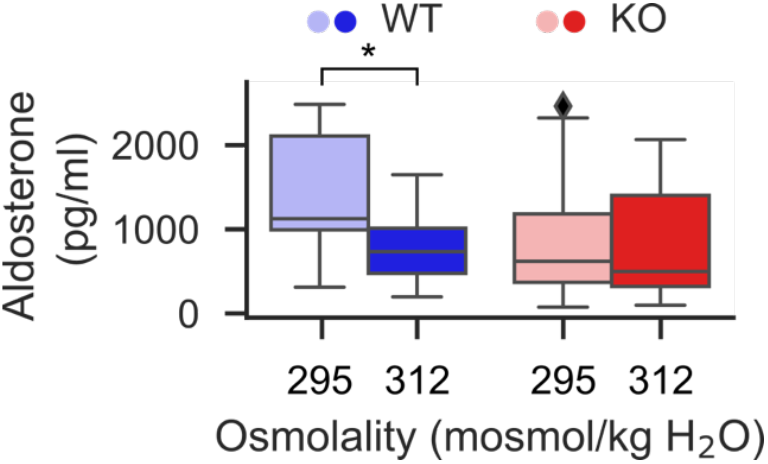
Incubation of whole adrenal glands in vitro demonstrates osmolality-dependent aldosterone secretion in WT mice. They showed an increase in aldosterone production when exposed to hypoosmolality (295 mosmol/kg H_2_O) for 120 minutes. This effect was absent in KO mice (median: WT @295 mosmol/kg H_2_O: 1125 pg/ml, @312 mosmol/kg H_2_O: 734 pg/ml, n = 12, T = 10, p = 0.02, *; KO @295 mosmol/kg H_2_O: 619 pg/ml, @312 mosmol/kg H_2_O: 499 pg/ml, n = 15, T = 59, p = 0.98; Wilcoxon signed-rank test). Box plots are shown as defined in Materials & Methods.

We confirmed that hypoosmolar conditions increased aldosterone production in WT but not KO mice (Fig. 3). Neither genotype showed changes when the extracellular osmolality was held constant at 312 mosmol/kg H_2_O (Supplementary Fig. 1).

### *Clcn2* KO mice exhibit increased calcium spiking under hypoosmotic conditions

Gain-of-function mutations in ClC-2 increase ZG intracellular calcium, presumably via a constant depolarization and subsequent increase in spiking activity^24,25^. To study whether the observed changes in intracellular calcium concentrations in our knock-out model were the result of alterations of the calcium spiking pattern, we recorded calcium signals with a higher temporal resolution using Calbryte 520 AM. In samples from both genotypes, the typical oscillatory changes in intracellular calcium were visible (Fig. 4A), with small changes to these patterns as indicated by higher frequencies and longer bursts in the KO (Supplementary Fig. 2).

**Figure 4.**
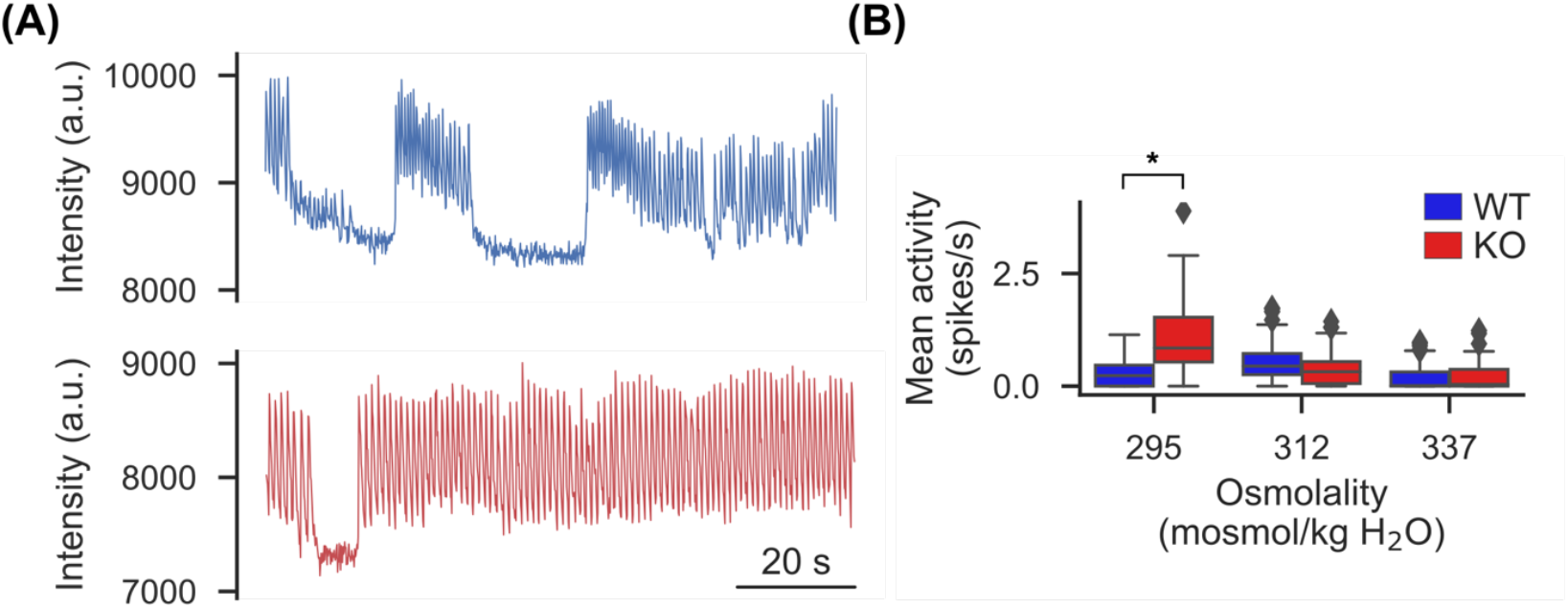
Exposure to hypoosmotic extracellular solutions results in higher calcium spiking activity in *Clcn2* KO mice. **(A)** Representative recordings of individual ZG cells in an acute slice preparation from a WT (blue, top) and *Clcn2* KO (red, bottom) mouse. These recordings were performed at 295 mosmol/kg H_2_O. **(B)** Ca^2+^ spiking activity (spikes per second) recorded in ZG cells over a period of 10 minutes. Activity was higher in KO than in WT mice under hypotonicity (WT (295/312/337): n= 5/5/4 recordings, KO (295/312/337): n =5/5/5 recordings; p (295/312/337) = 0.05(*)/0.09/0.98; likelihood ratio test of linear mixed models; further information can be found in Supplementary Table 3).

Given the effect of hypoosmolality on ClC-2 activation, we compared calcium spiking under different conditions: While there were no apparent differences between genotypes under prolonged (>90 minutes) isotonic (312 mosmol/kg H_2_O) and hypertonic (337 mosmol/kg H_2_O) conditions, hypotonic exposure (295 mosmol/kg H_2_O) resulted in significantly higher spiking frequency in KO mice (Fig. 4B).

### Calcium spiking frequencies are increased in ClC-2 KO mice

Following an acute change of extracellular osmolality, both WT and KO mice showed an immediate rise in calcium activity (Fig. 5A). In WT, activity then decreased to a low steady state while the activity in KO mice remained elevated in comparison (Fig. 5A).

**Figure 5.**
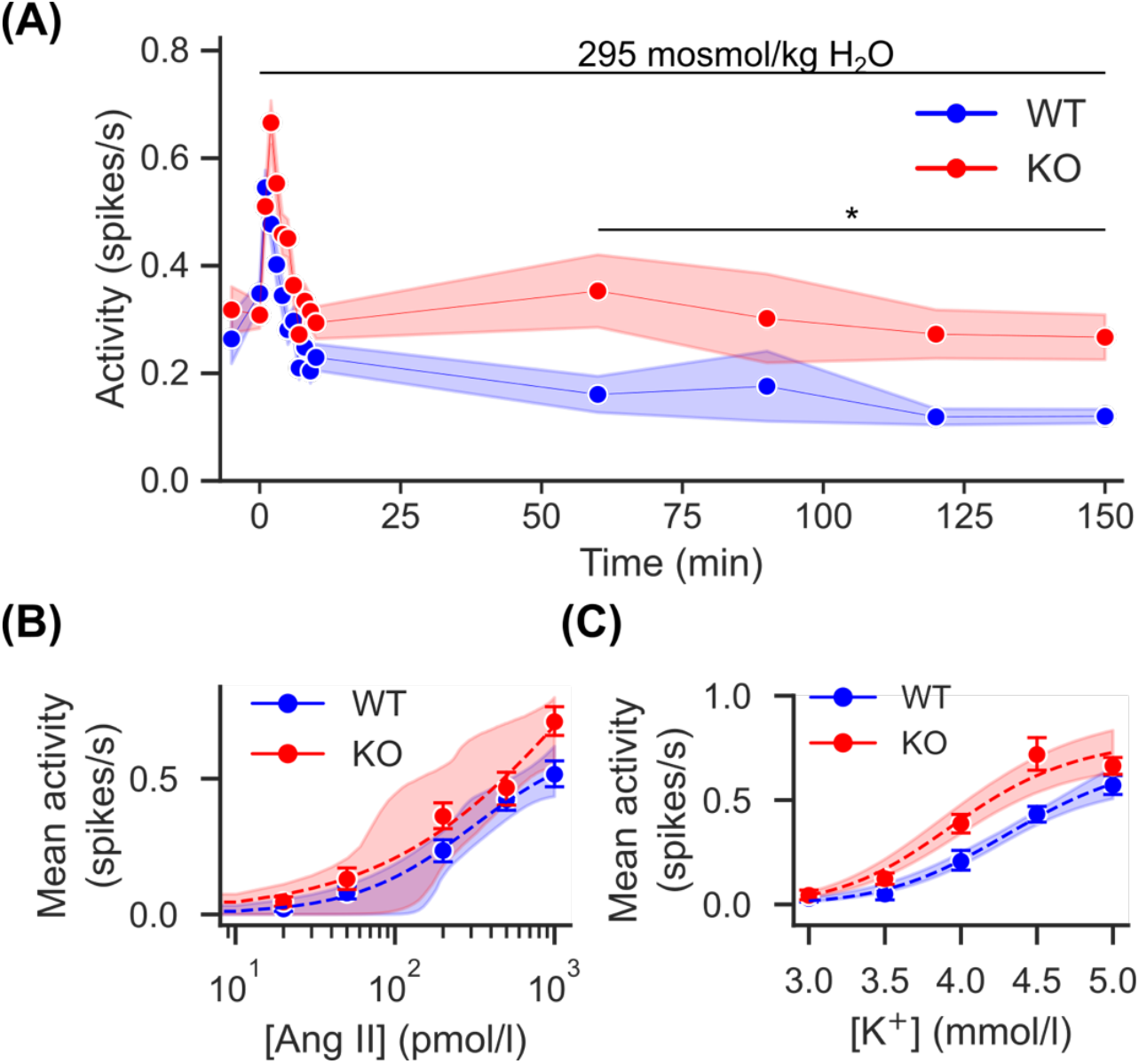
ZG cells from *Clcn2* KO mice show different adaptation of calcium signaling when exposed to hypoosmolality. **(A)** Changes in the spiking activity upon exposure to hypoosmolality starting from isotonic conditions (n_WT_ = 6 animals; n_KO_ = 6 animals). Immediately after the switch to a hypotonic environment (starting at t = 0 min), spiking activity rose in both genotypes. The slightly higher mean activity observed in KO mice during the first 10 minutes, which likely correspond to the RVD, was not significantly different to WT (p = 0.081, χ^2^(1) = 3.25). During longer incubation, however, the activity in KO remained significantly elevated relative to WT cells (p = 0.009, χ^2^(1) = 6.91). Statistical analysis was performed using likelihood ratio tests of linear mixed models. **(B)** Calcium spiking activity increases with extracellular Ang II concentrations. There is no significant difference between ZG cells from WT and *Clcn2* KO mice at an osmolality of 295 mosmol/kg H_2_O. Dashed lines represent a fit with a Hill equation, the shaded areas comprise the 95% confidence intervals of all combined fits. Measured data is shown as mean (circles) with 95% CI of the measured data shown as errorbars. Fitting of the Hill equation was performed using a bootstrap algorithm with 10,000 resamples. WT (mean and 95% CI): V_max_ 0.65 (0.46-2.28), K_50_=311.31 (150.51-4626.54), n=1.16 (0.71-10.0); KO: V_max_ 1.59 (0.54-85.55), K_50_=1435.79 (65.35-2116015.40), n=0.71 (0.55-10.0). **(C)** The potassium dependence of calcium spiking activity is shifted towards lower concentrations at an osmolality of 295 mosmol/kg H_2_O. Dashed lines represent a fit with a Hill equation, the shaded areas comprise the 95% confidence intervals of all combined fits. Measured data is shown as mean (circles) with 95% CI of the measured data shown as errorbars. Fitting of the Hill equation was performed using a bootstrap algorithm with 10,000 resamples. WT (mean and 95% CI): V_max_ 0.73 (0.60-1.08), K_50_=4.37 (4.18-4.80), n=10.0 (7.47 −10.0); KO: V_max_ 0.81 (0.69-0.94), K_50_=3.99 (3.84-4.15), n=10.0 (9.31-10.0).

Closer analysis showed unchanged Ang II-dependent spiking between genotypes (Fig. 5B; Supplementary Fig. 2) but revealed a leftward shift in potassium-dependent activation of calcium spiking in KO mice (Fig. 5C).

This shift manifested as both, an increased mean intra-burst frequency and longer burst durations (Supplementary Fig. 2). Taken together, these results suggest that lower calcium concentrations in the KO are not the result of lower spiking frequencies.

### Systemic hypoosmolality increases aldosterone production

To test whether ClC-2’s role in osmo-sensing affects aldosterone production in vivo, we induced chronic hypoosmolality in WT mice using desmopressin. This is a synthetic analog of vasopressin and activates V2 receptors in the kidney resulting in excessive water reabsorption, creating a hyponatremic hypoosmolality. For a continuous release, we surgically implanted osmo-pumps that slowly released desmopressin over several days.

Both genotypes showed significant hyponatremia (mean ± SD; WT: 112.3 ± 8.6 mM, n = 7 mice; KO: 117.2 ± 10.1 mM, n = 9 mice; p = 0.32, t = −1.03, Student’s t-test) in contrast to non-implanted WT mice from a previous study^27^ (mean ± SD; 146.6 ± 2.3 mM, n = 20, vs implanted WT: p = 2.15×10^−15^, t = −16.7; vs implanted KO: p = 4.5×10^−13^, t = −12.5; Student’s t-test with Bonferroni correction). Aldosterone values were markedly higher than those measured in non-implanted mice previously reported by our group^24^ (Fig. 6A).

**Figure 6.**
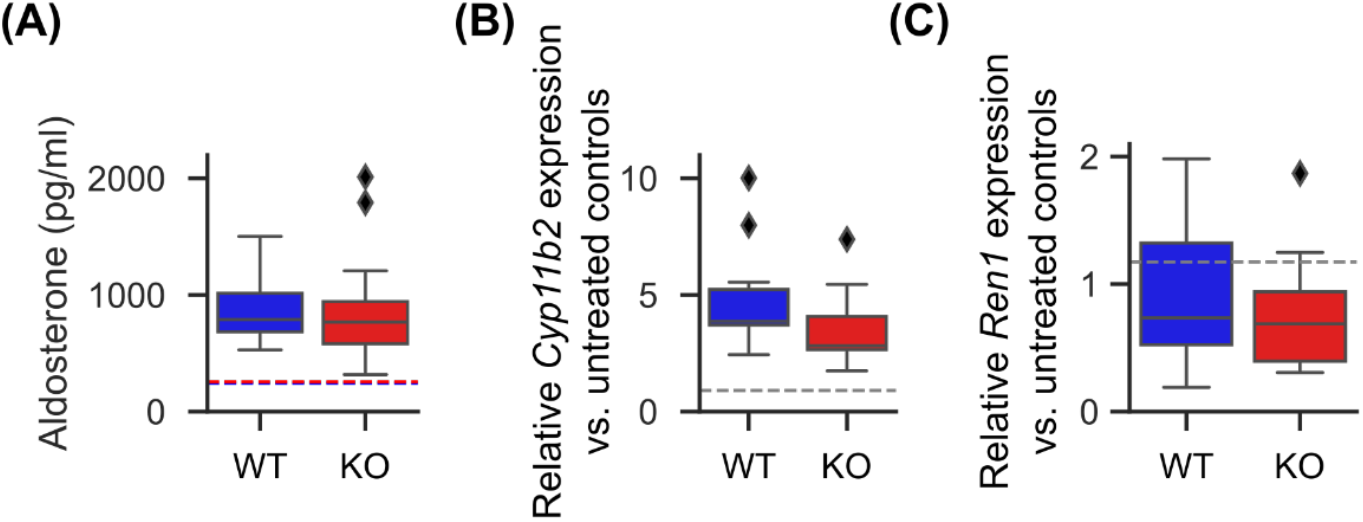
Aldosterone production under hyponatremic hypoosmolality is similarly increased in *Clcn2* KO compared to WT mice. **(A)** Mean serum aldosterone concentration after 1 week of forced hyponatremia. WT and KO values are similar (mean ± SD, WT: 904 ± 327 pg/ml, n = 10; KO: 886.74 ± 532 pg/ml, n = 12, p = 0.57, U = 69; Mann-Whitney-U test). Aldosterone concentrations are, however, higher than in untreated controls reported previously^24^ (WT, blue: 243 pg/ml; KO, red: 255 pg/ml; dashed lines). **(B)** The relative expression of aldosterone synthase (*Cyp11b2*) is significantly higher than in untreated controls (mean ± SD, untreated C57BL/6 from^27^: 0.90 ± 0.48 (mean shown as dashed line), n = 8; WT: 4.82 ± 2.28, n = 11, versus untreated: p < 0.001, U = 0; KO: 3.57± 1.63, n = 12; versus untreated: p < 0.001, U = 1; Mann-Whitney-U test). However, while the mean values suggest less *Cyp11b2* expression in KO, this difference was not statistically significant (KO versus WT: p = 0.069, U = 36; Mann-Whitney-U test). **(C)** The relative expression of Renin (*Ren1*) was similar in WT and KO adrenal glands compared to untreated controls (mean ± SD: untreated C57BL/6 from^27^ : 1.17 ± 0.86, n = 11 (mean is shown as dashed line); WT: 0.91 ± 0.58, n = 10, versus untreated: p = 0.46, U = 66; KO: 0.77 ± 0.45, n = 12; versus untreated: p = 0.281, U = 84; Mann-Whitney-U test). There was no statistical difference between implanted WT and KO mice (p = 0.668, U = 53; Mann-Whitney-U test).

The expression of aldosterone synthase (*Cyp11b2*) appeared slightly higher in WT over KO, but the difference was not statistically significant (Fig. 6B). Renin (*Ren1*) expression was highly variable with no appreciable difference between the two genotypes and no increase over untreated controls (Fig. 6C).

## Discussion

Studies using perfused whole canine adrenal glands have demonstrated that even small decreases in osmolality amplify aldosterone secretion^35^. This process was found to be chloride-dependent^32^, but the molecular mechanisms remained unclear.

We here link this mechanism to the ClC-2 chloride channel. The high levels of intracellular chloride in the ZG^9^ (Fig. 1) combined with the known hyperpolarized resting membrane potential^2^ suggest efflux of chloride from the ZG as a primary mechanism of ClC-2 upon its activation. Given the known osmo-sensitivity of this channel, we hypothesized that hypoosmolality may serve as the activating stimulus. In agreement with this, we observed lower chloride concentrations in the ZG of WT compared to *Clcn2* KO mice under sustained hypoosmolar conditions (Fig. 1).

Upon a hypoosmolar challenge, cells initially quickly extrude ions, particularly chloride, to counteract cell swelling in a process called rapid volume decrease (RVD)^36^. This phase likely corresponds to the sharp initial increase in calcium spiking seen upon hypoosmolar challenge (Fig. 5A), with activation of several ion conductances that mediate RVD^36,37^. ClC-2 apparently only mediates a small effect during this phase as seen by the small, non-significant difference between WT and KO (Fig. 5A).

Long-term osmotic adaptation requires the slower efflux of other, non-ionic substances (osmolytes), allowing for the re-entry of ions and restoration of ion balances^37^. This likely corresponds to the period after the initial peak, similar as in other cell types^38,39^. Here, ClC-2 is responsible for maintaining chloride efflux (Fig. 1) and the loss of ClC-2 leads to higher calcium spiking activity compared to WT (Fig. 5A). It may be that ClC-2 serves as a recycling pathway for other ion transporters during this phase, such as the highly expressed NKCC1^25^, ultimately facilitating the (re-)entry of Na^+^ and K^+40^.

A chloride efflux depolarizes ZG cells and activates voltage-gated calcium channels^24,25^, allowing for calcium influx. Accordingly, intracellular calcium concentrations rose in the ZG of WT when exposed to hypoosmolar conditions, an effect that was abrogated in the KO (Fig. 2). This was more pronounced for stimulation with Ang II rather than potassium, likely due to the additive contribution of intracellular stores to the response to Ang II^41^. The effect of K^+^ primarily depends on depolarization and influx via voltage-dependent calcium channels^42^, likely affecting the same pathway as hypoosmolality (Fig. 5).

In line with the known role of intracellular calcium in regulating aldosterone production^3,4^, aldosterone production of adrenal glands ex vivo rose upon hypoosmolar stimulation, an effect that was again attenuated by the ClC-2 knockout.

In vivo, the effect of hypoosmolality on aldosterone production does not solely depend on ClC-2. Upon administration of desmopressin, aldosterone levels increased in both WT and KO animals, as did levels of *Cyp11b2* expression, despite unchanged levels of the upstream regulator renin (*Ren1*; Fig. 6). Interestingly, both aldosterone and *Cyp11b2* levels were slightly, though not significantly, lower in KO animals. However, statistical power to detect this small effect was only about 40% (*post hoc* analysis), suggesting that larger sample sizes would be required to further investigate the in vivo role of ClC-2 in hypoosmolality-induced aldosterone production. Of note, in vivo, hypoosmolality was achieved through hyponatremia, unlike the situation ex vivo, in which sodium concentrations remained constant.

The higher frequency of calcium spiking in KO animals under hypoosmolar conditions remains unexplained (Fig. 4 and 5) but suggests that baseline and/or peak calcium concentrations must be lower in KO animals (Fig. 2). Interestingly, mice carrying a gain-of-function mutation in the Ca_V_3.2 T-type voltage-gated calcium channel^26^ show the reverse phenotype, with higher intracellular calcium concentrations despite slower calcium spiking. These effects may be linked to calcium-dependent potassium channels in the ZG^43^ and a more robust hyperpolarization, delaying the onset of oscillatory depolarizations in the presence of high calcium concentrations.

While our study was conducted in mice, ClC-2 is also expressed in human adrenal glands^9^ and may have similar functions there. Our slice-based approach cannot capture all aspects of integrated organ function as demonstrated in perfused whole-gland studies^31^, yet it enables detailed investigation of cellular mechanisms and ion fluxes that would be difficult to measure in intact organs.

In summary, we have found a role of ClC-2 mediated chloride efflux in the maintenance of chloride homeostasis in the ZG in response to hypoosmolality. This may at least partially explain the known chloride dependence of the hypoosmolality-induced aldosterone synthesis.

## Methods

### Breeding *of Clcn2* KO mice

A single ClC-2-deficient (*Clcn2* KO) mouse was originally provided by Thomas Jentsch (FMP Berlin)^20^. Breeding was performed in the Forschungseinrichtungen für Experimentelle Medizin at Charité – Universitätsmedizin Berlin. Heterozygous mice were generated by backcrossing with C57BL/6NCrl mice. All mice were housed in a 12h light/dark cycle (6 a.m to 6 p.m) with unrestricted access to food and water, environmental enrichment and under specific-pathogen free conditions.

### Implantation of osmotic pumps filled with desmopressin acetate

First, osmotic pumps (ALZET, Model 1004) were filled with desmopressin acetate (0.002 mg/ml Minirin prepared in sterile 0.9% NaCl) under a sterile workbench according to the manufacturer’s instructions. Before implantation, the filled pumps were activated in sterile 0.9% NaCl solution at 37°C 48h.

Mice were anesthesized using isoflurane (induction chamber: oxygen flow of 1 l/min 100 % oxygen, 5 Vol.-% isoflurane; continuous anesthesia under an inhalation mask: oxygen flow of 1 l/min 100 % oxygen, 2-3 Vol.-% isoflurane on top of a heating blanket (37 °C)). Carprofen (5 mg/kg) was injected subcutaneously prior to the procedure for analgesia.

A skin incision of about 0.5-1 cm was made between the scapulae, and a small pocket was formed using blunt scissors. The pumps were inserted with the flow moderator pointing away from the incision. The incision was then closed with a wound closure system (autoclip), and each mouse was monitored until it was fully awake.

### Induction of hypoosmolality in mice

After implantation, mice were housed in regular home cages for 5 days. Water and food (AIN76A diet gel, ClearH_2_O; exchanged daily) were provided ad libitum.

Afterwards, blood, urine and organs were collected following i.p. anesthesia (ketamine, 100 mg/kg and xylazine, 10 mg/kg) and terminal blood collection via cardiac puncture. To determine renin concentrations, the first 100 µl of blood were collected into tubes containing 10 µL EDTA (0.1 M). The remaining blood was collected into Microtainer EDTA tubes (Becton Dickinson) and Lithium-Heparin tubes (Becton Dickinson). All blood samples were centrifuged (10 minutes, 2000 g, 4 °C), and the supernatant (plasma) was stored at −20 °C until further analysis.

The aldosterone plasma levels were analyzed using an ELISA (RE52301, IBL International GmbH). Blood plasma sodium was measured using an AU480 Clinical Chemistry Analyzer (Beckman Coulter) at the Animal Phenotyping Facility at the Max-Delbrück-Centrum Berlin.

### Aldosterone production of explanted adrenal glands

The explanted adrenal glands were carefully cut by hand into four equal-sized pieces and stored in carbogen-gassed BBS_Supp_ at 312 mosmol/l kg H_2_O supplemented with 2 mmol/l CaCl_2_ (Table 1) in a cell culture insert in a 12-well plate (pore diameter: 1 µm, Greiner Bio-One). The well and the inside of the cell insert were each filled with 1000 μl BBS ^+^ (Table 1), supplemented with 100 pmol/l Ang II and continuously gassed with carbogen at RT. After 30 minutes, 500 µl of the solution (outside of the insert) was collected (sample t_0_). Since two adrenal glands were collected from each animal, the solutions were exchanged in one well and insert with fresh BBS ^+^ and in the other with BBS ^+^ solution, supplemented with 100 pmol/l Ang II. After 90 minutes, solutions were again exchanged for fresh solutions of the same osmolality and 500 µl were collected after another 30 minutes. The aldosterone concentrations were measured using ELISA (RE52301, IBL International GmbH) according to the manufacturer’s instructions.

### Slice preparation

Mice were anesthetized by using isoflurane (400 μl as an open drop in a 2 l beaker) and subsequently euthanized by cervical dislocation. Both adrenal glands were removed from the mice via the abdominal cavity. Any remaining fat surrounding the adrenal glands was carefully removed. Afterwards, the adrenal glands were transferred into ice-cold bicarbonate-buffered solution (BBS_295_), which was continuously gassed with carbogen (95% O_2_ + 5% CO_2_).

Before the adrenal glands were cut in a vibratome at 120 µm thickness (7000 smz-2, Campden Instruments), they were embedded in 4% low-melting agarose. During the slicing process, the organ and slices were stored in ice-cold BBS_295_ solution, continuously gassed with carbogen. Afterwards, slices were transferred into a carbogen gassed BBS_295_ at 35 °C for 15 minutes and then stored for up to 6 h at RT in carbogen gassed BBS_295_ supplemented with 2 mmol/l CaCl_2_ (BBS_Supp,295_).

### Preparation of diH-MEQ

For Cl^−^ imaging experiments in living cells, MEQ was reduced to the cell-permeable form diH-MEQ. This synthesis was performed according to the manufacturer’s instructions. All steps were performed while purging oxygen from the reaction using a slow stream of nitrogen. In detail, diH-MEQ was prepared from 5 mg of MEQ (21250, AAT Bioquest) and dissolved in 0.1 ml of distilled water. During the dissolution processes, a small amount of (20-50 µl) of a 12 % (120 mg/ml) aqueous solution of sodium borohydride (424270010, Thermo Scientific Chemicals) was prepared for immediate use. 10 µl (32 µmol) of the sodium borohydride solution was slowly added. After 30 minutes, the yellow diH-MEQ was extracted from the solution with anhydrous ethyl acetate (270989-250ML, Sigma-Aldrich). The upper organic layer containing diH-MEQ was transferred into a 4 ml sample vial (WHEATON^®^ amber with cap, Merck). The extraction step was repeated twice with an additional 0.5 ml of ethyl acetate each time. After the organic extracts were combined, 150 mg anhydrous sodium sulfate (S6547-500G, Sigma-Aldrich) was added to the vial and gently mixed to dry the organic layers and transferred to a 5 ml Schlenk sample tube (SP Wilmad-LabGlass, VWR). The organic solvent was now carefully evaporated under vacuum in a water bath at 35-45 °C. The solid diH-MEQ (yellowish-light green oil) was dissolved in 100 µl DMSO/10 % Pluronic F-127 (∼150 mmol/l) by vortexing thoroughly for 30 seconds. 2 µl aliquots were prepared and transferred into 1.5 ml screw cap micro tubes that were tightly sealed under N_2_ and stored at −80 °C for use within 1 week.

### Staining of adrenal slices with Calbryte 520 AM, Fura-2 AM or diH-MEQ

For staining the slices, a cell culture insert (pore diameter: 1 µm, Greiner Bio-One) was added to a single well of a 24-well plate. The well (outside of the insert) was filled with 750 μl BBS_2K_ ^+^_295_ and continuously gassed with carbogen. The insert was filled with 250 μl of BBS_2K_ ^+^_295_ containing initially either 64 μmol/l Fura-2 AM (21023, AAT Bioquest; final well concentration: 16 μmol/l) and 0.16% Pluronic F-127 or 37 μmol/l Calbryte 520 AM (20650, AAT Bioquest; final well concentration: 9.25 μmol/l) and 0.001% Pluronic F-127. For diH-MEQ, the insert was filled with 250 μl containing initially either 600 µmol/l diH-MEQ (final well concentration: 150 μmol/l) and 0.4 % Pluronic F-127. Regardless of the dye used, slices were left in the dyeing chamber for 1h at RT in the dark.

### Measurement of Ca^2+^ signals using Calbryte 520 AM

Calbryte 520 stained slices were transferred into the recoding chamber of the microscope (Scientifica SliceScope). The stained slices in the recoding chamber were continuously perfused with carbogen-gassed solutions, heated by an inline heating coil to 30 ± 1 °C. For Calbryte 520 measurements, a blue LED (channel 2B, pE-300 ultra, CoolLED) passing through a 474/27 nm bandpass filter (AHF Analysentechnik) was used to excite the dye. Signal emissions were collected by a 40 × /0.8 NA objective (LUMPlanFL N, Olympus), filtered through a 432/523/702 nm triple-band filter (AHF Analysentechnik) and recorded using a sCMOS camera (OptiMOS, QImaging) every 100 ms with a 10 ms exposure.

### Measurement of absolute [Ca^2+^]_*i*_ using Fura-2 AM

Fura-2 stained slices were transferred into the recoding chamber of the microscope (Scientifica SliceScope). The stained slices in the recoding chamber were continuously perfused with carbogen-gassed solutions. For Fura-2 measurements, a light source with 340 and 385 nm LEDs (FuraLED, Cairn Research) was used. Emission light was collected using a 40 × /0.8 NA objective ((LUMPlanFL N, Olympus), a 510/84 nm bandpass filter (AHF Analysentechnik) and a camera (OptiMOS, QImaging). Every 100 ms, two images with 10 ms exposure each were taken during excitation with the 340 and 385 nm LED, respectively.

The calibration of Fura-2 signals was performed as previously described, using a calibration kit (Calcium Calibration Buffer Kit, Thermo Scientific), to obtain the 340/385 nm ratios to the corresponding [Ca^2+^]_*i*_ in Fura-2 stained adrenal gland slices.

### Measurement of Cl^−^ concentrations using diH-MEQ

diH-MEQ stained slices were transferred into the recording chamber of the microscope (Axioskop 2 FS, Zeiss). Slices were kept in the perfused chamber for 15 minutes before the start of the first recording to ensure an even distribution of the MEQ dye.

For MEQ measurements, a 340 nm LED (Thorlabs) passing through a 340/26 nm bandpass filter (Edmund Optics) was used for excitation. Emissions were recorded using a 40×/0.75 NA objective (Achroplan, Zeiss), a 510/84 bandpass filter (Edmund Optics) and a camera (GS3-U3-51S5M-C, FLIR) every 20s with a 50 ms exposure.

Two different approaches were used to calibrate the MEQ signals to absolute Cl^−^ concentrations: Using several different concentrations of Cl^−^ in a separate series of recordings (various-slice calibration) or after each measurement directly on the same slice (same-slice calibration) using only two Cl^−^ concentrations.

In both cases, intracellular Cl^−^ concentrations were clamped to defined values (0-80 mmol/l Cl^−^) by incubation with standard solutions (BBS_5K_ ^+^_312_ or BBS_5K_ ^+^_295_) supplemented with 10 µmol/l nigericin (11437, Cayman Chemicals) and 20 µmol/l tributyltin chloride (T50202-100G, Sigma-Aldrich).

### Analysis of fluorescence microscopy

Microscopy recordings were taken using Micro-Manager 2.0^44^ and processed using Fiji^45^. After manual selection of individual cells and export from Fiji, further analysis was performed using custom python scripts^3,26^.

### Quantitative real-time PCR

The extraction/isolation of total RNA from adrenal glands and kidneys stored RNAlater, (Sigma-Aldrich) at −20 °C was performed using the RNeasy Mini Kit (Qiagen). Reverse transcription was performed using Quantitect RT Kit (Qiagen). For adrenal and kidney cDNA samples, the TaqMan real-time PCR expression master mix (Applied Biosystems) was used with *Gapdh* (Mm99999915g1, housekeeping gene), *Cyp11b2* (Mm00515624m1) and *Ren1* (Mm02342887_mH) probes. The gene expression was evaluated relative to *Gapdh*, and mean ΔCT of WT controls and expressed as 2^ΔΔCt^ (fold change).

### Statistics

Data were statistically analyzed using custom Python and R scripts. In Python, the Scipy^46^ library was used for analysis. In particular, the scipy.stats.shapiro function was used to assess normality. If normality was violated, the scipy.stats.mannwhitneyu or scipy.stats.wilcoxon functions were used for statistical comparisons, otherwise the scipy.stats.ttest_ind function was used as stated in the text. Calcium concentrations were analyzed using linear mixed models in R using the lme4 package^47^. All box plots shown follow Tukey style: the box encloses by the 25^th^ to 75^th^ percentile with the central line defining the median. Whiskers are 1.5 × inter-quartile range (IQR) with outliers shown as diamonds.

## Data availability

Raw microscopy video files can only be provided upon reasonable request and must be individually arranged due to their file sizes. All exctracted data and scripts used to generate the analyses and figures presented within this manuscript are available at zenodo^48^ (https://zenodo.org/records/15051779).

## Author contributions

**Marina Volkert:** Development of methodology, acquisition of data, analysis of data, preparation of figures, editing the manuscript. **Hoang An Dinh:** Acquisition of data, editing the manuscript. **Ute I. Scholl:** Conceptualization, preparation of figures, writing and editing the manuscript. **Gabriel Stölting:** Conceptualization, analysis of data, curation of data, supervision, writing and editing the manuscript.

## Acknowledgements

We would like to thank Sarah Döring, Nico Brüssow, Marie Cotta and Ana Lucia Huitron Carrizales for mouse genotyping. This research was funded by the Deutsche Forschungsgemeinschaft (STO 1260/1-1 for GS) and Stiftung Charité (BIH_PRO_406 to UIS).

