## Supplementary Material for "ClC-2 contributes to hypotonicity-induced adrenal aldosterone secretion"

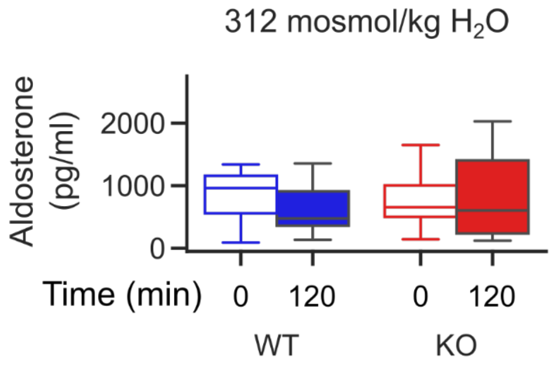


**Supplementary Figure 1.** Incubation of whole adrenal glands in vitro shows no significant changes in aldosterone production over 2 hours. After removal of adrenal glands from mice, they were individually incubated in solution containing 5 mmol/l $K^+$ and 100 pmol/l Ang II at isoosmotic 312 mosmol/kg $H_2O$. After 30 minutes, the supernatant was collected and replaced with fresh solution with the same concentrations of $K^+$ and Ang II and osmolality. After another 90 minutes, these were replaced with fresh solutions of the same composition. 30 minutes later, the supernatant was again collected. Aldosterone concentrations at t=0 min (open boxes) and t=120 min (filled boxes) were determined using ELISA. There was no significant change in aldosterone production after 120 minutes (WT 0 min: 961 pg/ml, WT 120 min: 477 pg/ml, n = 11, T = 15, p = 0.12; KO 0 min: 655 pg/ml, KO 120 min: 605 pg/ml, n = 13, T = 36, p = 0.54; Wilcoxon signed-rank test).

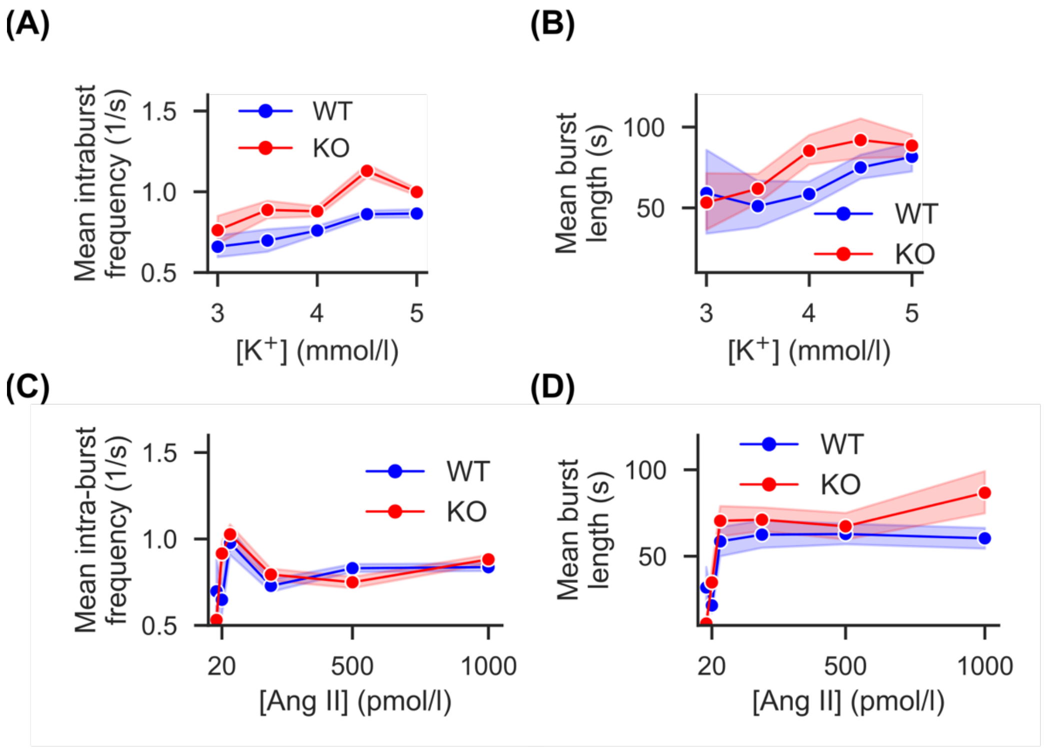


**Supplementary Figure 2.** *Clcn2* KO mice exhibit potassium- but not angiotensin II-dependent alterations to calcium spiking at 295 mosmol/kg $H_2O$.

**(A)** The intra-burst frequency shows modest dependence on extracellular potassium. Data are plotted as mean values (circles) ± SD (shaded areas).

**(B)** The mean length of bursts is also dependent on extracellular potassium. Data are plotted as mean values (circles) ± SD (shaded areas).

**(C)** Intra-burst frequencies are similar, independent of the applied concentration of angiotensin II. Data are plotted as mean values (circles) ± SD (shaded areas).

**(D)** Burst length is not different between genotypes, irrespective of the concentration of angiotensin II except for very high values. Data are plotted as mean values (circles) ± SD (shaded areas).

**Supplementary Table 1: Statistical information for the angiotensin II dependent calcium concentrations in ZG cells (Fig. 2A)**

| | | | | $[Ca^{2+}]$ (nmol/l) | | | |
|---|---|---|---|---|---|---|---|
| **Genotype** | **Osmolality (mosmol/kg $H_2O$)** | **[Ang II] (pmol/l)** | **N (cells/ female mice/ male mice)** | **Mean** | **s.d.** | **P vs. 312** | **$\chi^2$ df=1** |
| **WT** | **295** | **20** | 169/2/2 | 903.04 | 157.54 | 0.02 | 5.08 |
| | | **100** | 169/2/2 | 911.85 | 148.25 | 0.03 | 4.98 |
| | | **500** | 169/2/2 | 1004.54 | 150.74 | 0.01 | 6.56 |
| | **312** | **20** | 148/3/4 | 826.83 | 225.02 | - | |
| | | **100** | 148/3/4 | 837.16 | 192.26 | - | |
| | | **500** | 148/3/4 | 913.92 | 181.08 | - | |
| **KO** | **295** | **20** | 134/3/3 | 846.15 | 185.54 | 0.59 | 0.29 |
| | | **100** | 134/3/3 | 880.43 | 165.54 | 0.51 | 0.44 |
| | | **500** | 134/3/3 | 960.37 | 163.50 | 0.58 | 0.30 |
| | **312** | **20** | 90/3/4 | 846.93 | 143.05 | - | |
| | | **100** | 90/3/4 | 856.18 | 144.09 | - | |
| | | **500** | 90/3/4 | 930.15 | 133.25 | - | |

**Supplementary Table 2: Statistical information for the potassium dependent calcium concentrations in ZG cells (Fig. 2B)**

| | | | | $[Ca^{2+}]$ (nmol/l) | | | |
|---|---|---|---|---|---|---|---|
| **Genotype** | **Osmolality (mosmol/kg $H_2O$)** | **$[K^+]$ (mmol/l)** | **N (cells/ female mice/ male mice)** | **Mean** | **s.d.** | **P vs. 312** | **$\chi^2$ df=1** |
| **WT** | **295** | **3** | 182/2/2 | 915.76 | 188.08 | 0.03 | 4.68 |
| | | **4** | 182/2/2 | 1029.38 | 202.05 | 0.07 | 3.22 |
| | | **5** | 182/2/2 | 1096.04 | 191.20 | 0.44 | 0.60 |
| | **312** | **3** | 109/3/4 | 825.79 | 160.90 | - | |
| | | **4** | 109/3/4 | 939.18 | 195.14 | - | |
| | | **5** | 109/3/4 | 1082.25 | 209.96 | - | |
| **KO** | **295** | **3** | 133/3/2 | 842.68 | 208.73 | 0.98 | $8\times10^{-4}$ |
| | | **4** | 133/3/2 | 932.02 | 198.88 | 0.82 | 0.05 |
| | | **5** | 133/3/2 | 980.86 | 178.63 | 0.66 | 0.20 |
| | **312** | **3** | 103/2/3 | 840.36 | 160.37 | - | |
| | | **4** | 103/2/3 | 929.59 | 181.68 | - | |
| | | **5** | 103/2/3 | 1004.90 | 161.45 | - | |

**Supplementary Table 3: Statistical information for the osmolality-dependent spiking activity in ZG cells (Fig. 4B)**

| | | | Mean activity (spikes/s) | | | |
|---|---|---|---|---|---|---|
| **Genotype** | **Osmolality (mosmol/kg $H_2O$)** | **N (cells/ female mice/ male mice)** | **Mean** | **s.d.** | **P WT vs. KO** | **$\chi^2$ df=1** |
| **WT** | **295** | 108/2/3 | 0.32 | 0.33 | 0.05 | 3.87 |
| | **312** | 284/2/3 | 0.49 | 0.40 | 0.09 | 2.84 |
| | **337** | 157/2/2 | 0.17 | 0.25 | 0.98 | $9\times10^{-4}$ |
| **KO** | **295** | 141/3/2 | 0.90 | 0.78 | | |
| | **312** | 245/1/4 | 0.37 | 0.33 | | |
| | **337** | 144/3/2 | 0.22 | 0.27 | | |
